# Trait Resilience Modulates the Association Between Cortisol and Aperiodic Neural Dynamics

**DOI:** 10.64898/2026.07.09.737399

**Authors:** Kar Fye Alvin Lee, Suhail T. Asharaf, Li Liang, Tatia M.C. Lee

**Affiliations:** Laboratory of Neuropsychology and Human Neuroscience, Department of Psychology, The University of Hong Kong, Hong Kong, China; InnoCentre of Clinical Neuropsychology, The University of Hong Kong, Hong Kong, China

## Abstract

Cortisol, our stress hormone, exerts widespread influence on neural activity. However, its influence on the aperiodic component of the electroencephalography power spectrum remains to be investigated. Given individual differences in the capacity to cope with stress and adversity, it also remains unclear whether trait resilience moderates this relationship. Hence, the present study examined whether individual differences in trait resilience moderates the association between resting cortisol and aperiodic activity. Participants (*N*=145) completed various self-report questionnaires (e.g., trait resilience). Electroencephalography was recorded over a 20-minute baseline period, followed by salivary cortisol collection. The results revealed a significant moderating effect of trait resilience in the occipital scalp region. Specifically, higher cortisol concentration was associated with flatter 1/*f* slopes amongst individuals with low trait resilience, whereas this association was reversed amongst those with high trait resilience. Overall, our findings highlight the role of individual differences in trait resilience in shaping hypothalamic-pituitary-adrenal axis-related neural dynamics.

## Introduction

Our stress regulatory system plays an essential role in the physiological adaptation to environmental demands (McEwen, 2007), enabling us to navigate and thrive in a dynamic and complex world. Worryingly, chronic dysregulation of this system has been linked to poorer physical and mental health outcomes (Guidi et al., 2021; Jiang et al., 2025; Lee et al., 2025; Turner et al., 2020). Stress regulation involves the coordinated activity of multiple distinct neurophysiological pathways, including the sympatho-adrenal medullary (SAM) axis, hypothalamic-pituitary-adrenal (HPA) axis, and various frontotemporal cortical regions (Russell & Lightman, 2019; Ulrich-Lai & Herman, 2009; William Tank & Lee Wong, 2014). The HPA axis, in particular, facilitates sustained energy mobilisation through the release of cortisol from the adrenal cortex into the bloodstream, supporting prolonged physiological adaptations (Herman et al., 2016). Notably, elevated cortisol levels have widespread effects across various neural regions (Harrewijn et al., 2020). These effects may manifest as observable alterations in oscillatory neural dynamics, as evidenced by the robust association between attenuation of alpha activity and increased psychosocial stress (Vanhollebeke et al., 2022). Beyond oscillatory dynamics, there is a growing appreciation for the aperiodic activity of the electroencephalography (EEG) power spectrum as a distinct neural component (Donoghue et al., 2020), which remains underexplored in the context of our stress regulatory system. Therefore, examining aperiodic neural dynamics in relation to cortisol may enhance our understanding of the neural processes underlying HPA axis functioning and how their dysregulation contributes to adverse health outcomes.

The aperiodic component refers to the non-oscillatory activity of the EEG power spectrum, characterised by a power-law decrease in power as a function of increasing frequency (Miller et al., 2009). To date, there are several neurophysiological functional interpretations of this component. For instance, a longstanding neurophysiological account proposes that aperiodic activity is a non-specific proxy index for the large-scale neural population dynamics relating to excitatory-inhibitory balance, with steeper decay in power as a function of increasing frequency reflecting greater inhibitory over excitatory activity (Ahmad et al., 2022; Donoghue, 2025; R. Gao et al., 2017). Conversely, the neural gain control account views the decay in power as reflecting variations in neural noise, with steeper decay indicating lower levels of noise (Pertermann et al., 2019; Voytek et al., 2015). Regardless of the multiple competing interpretations, recent research using pupillometry as well as methylphenidate has shown that aperiodic activity is modulated by noradrenergic activity (Y. Gao et al., 2024; Pertermann et al., 2019). Importantly, this noradrenergic signalling is also modulated by the SAM axis, the fast-reacting pathway of the stress regulatory system. However, the potential influence of the HPA axis, the slower-reacting pathway, on aperiodic activity has yet to be studied. This gap is notable given that cortisol is the primary endocrine output of the HPA axis, which exerts downstream modulatory effects on GABAergic and glutamatergic neurotransmission (Mody & Maguire, 2012; Popoli et al., 2012), providing multiple plausible pathways through which HPA axis activity may influence aperiodic neural dynamics.

While moderate levels of stress may promote functional adaptation, chronic exposure to high levels of stress has been linked to alterations in the HPA axis functioning, which in turn increases the risk of negative mental health outcomes (Guidi et al., 2021; Lee et al., 2025). This cumulative physiological burden resulting from the chronic overloading of the stress regulatory system is referred to as allostatic load (McEwen, 1998, 2003, 2007; McEwen & Stellar, 1993). However, considerable individual differences exist in physiological responses associated with allostatic load, as evidenced by large variability in cortisol responses (Kudielka et al., 2009) and oscillatory EEG activity (Vanhollebeke et al., 2022). Such pronounced variance suggests that other unaccounted-for factors are at play. Notably, physiological response and behavioural outcomes following stress exposure differ as a function of resilience (Ebner & Singewald, 2017; Franklin et al., 2012), suggesting that resilience may serve as a potential moderator. Trait resilience, in particular, broadly refers to the capacity of an individual to cope with or adjust to stress and adversity (Connor & Davidson, 2003; Hu et al., 2015; Ong et al., 2006). Prior meta-analysis revealed a robust relationship between higher trait resilience and better mental health outcomes, with this relationship strengthening amongst individuals who have experienced, or were experiencing, adversity (Hu et al., 2015). Individuals with higher trait resilience has been found to be associated with more adaptive emotional regulation (Polizzi & Lynn, 2021), cognitive and affective flexibility (Genet & Siemer, 2011; Rademacher et al., 2023), and more efficient physiological stress responses (Kalisch et al., 2024). More recent research has also shown that dynamic shifts in aperiodic activity reflect adaptive metacontrol, the shift between flexibility-dominant and persistence-dominant control states depending on contextual demands, providing a neural account linking adaptive control states to coping styles and resilience (Yan et al., 2024, 2025). Indeed, recent research in adolescents found that aperiodic neural activity moderated the relationship between recent stress severity and risk of future depression recurrence, such that flatter aperiodic slopes were associated with a greater risk of future depression recurrence amongst individuals experiencing high levels of stress, but not low levels of stress (Schantell et al., 2026). However, despite potential relevance to HPA axis and metacontrol, it remains unclear whether individual differences in trait resilience, potentially serving as a psychological buffer, may modulate the relationship between cortisol and aperiodic neural dynamics.

The overarching aim of the present study was to examine the moderating effects of trait resilience on the potential relationship between HPA axis functioning and aperiodic neural dynamics. Building on previous research examining noradrenergic signalling and aperiodic activity (Y. Gao et al., 2024; Pertermann et al., 2019), our study focused on the slow-reacting pathway of the stress regulatory system (i.e., HPA axis) as indexed by resting-state cortisol concentration. In addition, we also accounted for the potential moderating effects of trait resilience to determine whether it acts as a psychological buffer to the effects of cortisol on aperiodic activity. Notably, a recent review highlighted multiple neurobiological potential pathways in which chronic stress exposure may drive hyperactivity in the HPA axis, triggering neuroinflammation and resulting in excitotoxicity and synaptic dysfunction (Sharma & Singh, 2026). Hence, we hypothesise that this HPA axis-driven shift toward heightened excitatory tone would be reflected as a flatter aperiodic slope of the EEG power spectrum. In addition, we also hypothesise that individuals with lower trait resilience will exhibit a more pronounced relationship between elevated cortisol levels and a flattened aperiodic slope. Conversely, we expect that higher trait resilience will serve as a protective buffer, decoupling this association and maintaining a more stable excitation-inhibition balance despite elevated cortisol levels.

## Methods and Materials

### Participants

The final sample for this study consisted of 145 adults with ages ranging from 18 to 44 years old (*Mage* = 24.28; *SDage* = 4.44; 44.14% males). The data were collected as part of a broader project on stress intervention (Lee et al., 2026). No power analysis was conducted for this study as the sample size was constrained by the design and data availability of the overarching project. Ethics approval was obtained from the Human Research Ethics Committee within the University (Reference Number: EA230490).

### Electroencephalography Acquisition, Preprocessing, Spectral Decomposition

Continuous EEG was recorded for 20 minutes to assess resting physiological activity in a naturalistic setting, in accordance with the 10/20 system at Fp1, Fp2, Fz, F4, F3, F8, F7, Cz, C4, C3, T4, T3, Pz, P4, P3, P8, P7, O2, and O1. Three additional electrodes were placed at AFz, M1, and M2, with M1 serving as the online reference and AFz as the ground. The signals were amplified and digitised at a resolution of 24-bit with a sampling rate of 500 Hz. The recorded EEG signals were processed offline using a zero-phase finite impulse response bandpass filter with a frequency range of 0.5–45 Hz. The signals were then re-referenced to a common average reference. The Generalized Eigenvalue De-Artifacting Instrument algorithm was used to attenuate non-neural activity from the EEG signals (Ros et al., 2025). For the spectral analysis, the EEG signal was first segmented into consecutive 1-minute epochs, after which the power spectral density (PSD) was computed for each segment. PSD was estimated using Welch’s method, with a 1000-sample Hann window, 50% overlap, mean averaging, constant detrending, and density scaling. The Fitting Oscillations and One Over *f* algorithm was used to fit the 1/*f* function on the PSD across 2–40 Hz to model the aperiodic activity (Donoghue et al., 2020). Thereafter, the residual PSD was calculated by subtracting the fitted 1/*f* aperiodic function from the raw PSD in log-space. Canonical oscillatory activity was then parameterised from the residual PSD by averaging within fixed frequency bands (i.e., delta: 1–4 Hz, theta: 4–8 Hz, alpha: 8–12 Hz, beta: 12–30 Hz) for each segment.

### Salivary Cortisol Assay

Salivary samples were collected using the Salivette^®^ Cortisol tubes in accordance with the manufacturer’s instructions, immediately after the baseline physiological recording. Salivary cortisol concentrations were quantified using liquid chromatography-tandem mass spectrometry (Raff & Phillips, 2019).

### Self-Report Questionnaires

The Connor-Davidson Resilience Scale (CD-RISC) (Connor & Davidson, 2003) was used to assess a general trait-like ability to cope with stress and adversity. Higher scores indicate greater perceived trait resilience. The WHO-5 Well-Being Index (WHO-5) (World Health Organization, 1998) was used to measure psychological well-being over the past two weeks. Higher scores indicate better mental well-being. The Groningen Sleep Quality Scale (GSQS) (Meijman et al., 1988) was used to assess subjective sleep quality of the previous night. Higher scores indicate poorer sleep quality.

### Statistical Analysis

All inferential statistical analyses were conducted using the Mixed Generalized Additive Model (GAM) Computation Vehicle with Automatic Smoothness Estimation package (Version: 1.9.1) in R (Wood, 2017). This framework allows for the flexible modelling of fixed and random effects (Hastie & Tibshirani, 1990). Given that there is empirical evidence supporting age and sex influence on EEG activity (Cave & Barry, 2021; De Blasio et al., 2025; Leroy et al., 2025), we included these covariates in all models. As diurnal cortisol is characterised by a gradual decrease throughout the day (Adam et al., 2017), cortisol sampling time was also included in the model as a covariate. Subject was included as a random effect to account for the hierarchical structure of the data (i.e., epochs and electrodes). Scaled-*t* distribution with identity link function was used.

All models were computed using restricted maximum likelihood estimation. The statistical analyses were performed in three stages. In the first stage, we evaluated the first hypothesis by examining the two-way interaction term (cortisol x electrode) to determine if the relationship between cortisol and 1/*f* exponent varied across the scalp. Thereafter, we evaluated the second hypothesis by including a three-way interaction term (cortisol x resilience x electrode). The second stage comprised a series of additional analyses testing the spectral specificity of the 1/*f* exponent findings by including the 1/*f* offset and canonical oscillatory activity as alternative outcome variables. In addition, if any of these spectral components exhibited similar interaction patterns, we conducted reciprocal models in which each measure was included as a covariate in the other model. The third stage comprised another series of analyses examining the specificity of the moderating effect of trait resilience by including psychosocial factors, including sleep quality and psychological well-being, as alternative moderators or covariates. All post hoc simple slope analyses were Holm-corrected.

## Results

### Descriptive Statistics

The descriptive statistics for the baseline cortisol concentration and various self-report questionnaires are reported in Table 1.

**Table 1.** Descriptive Statistics of the Baseline Cortisol and Self-report Questionnaires.

|  | *M* | *SD* | *Min* | *Max* |
| --- | --- | --- | --- | --- |
| Baseline Cortisol (ng/ml) | 1.16 | 1.01 | 0.12 | 6.42 |
| Trait Resilience | 36.23 | 5.65 | 19 | 50 |
| Psychological Well-being | 14.95 | 4.76 | 1 | 25 |
| Sleep Quality | 3.65 | 2.75 | 0 | 13 |

$N = 145$ . Trait Resilience, Psychological Well-being, and Sleep Quality were assessed through the Connor-Davidson Resilience Scale, WHO-5 Well-Being Index, and Groningen Sleep Quality Scale. Note that one participant did not complete the Groningen Sleep Quality Scale.

### Cortisol, Trait Resilience, and Aperiodic Activity

Regarding the first hypothesis, results revealed a significant cortisol x electrode interaction effect, χ*^2^*(18, 54918.14) = 233.50, *p* < .001. As shown in Figure 1, the association between cortisol concentration and 1/*f* exponent was strongest in the occipital, parietal, and frontocentral regions. By contrast, age (β = −0.008, *SE* = 0.005, *p* = .127), sex (β = −0.014, *SE* = 0.045, *p* = .753), and cortisol sampling time (β = −0.000, *SE* = 0.000, *p* = .238) were not significant predictors. However, epoch (β = 0.001, *SE* = 0.000, *p* < .001) was a significant predictor, such that the 1/*f* exponent increased in steepness over the baseline period.

**Figure 1.**
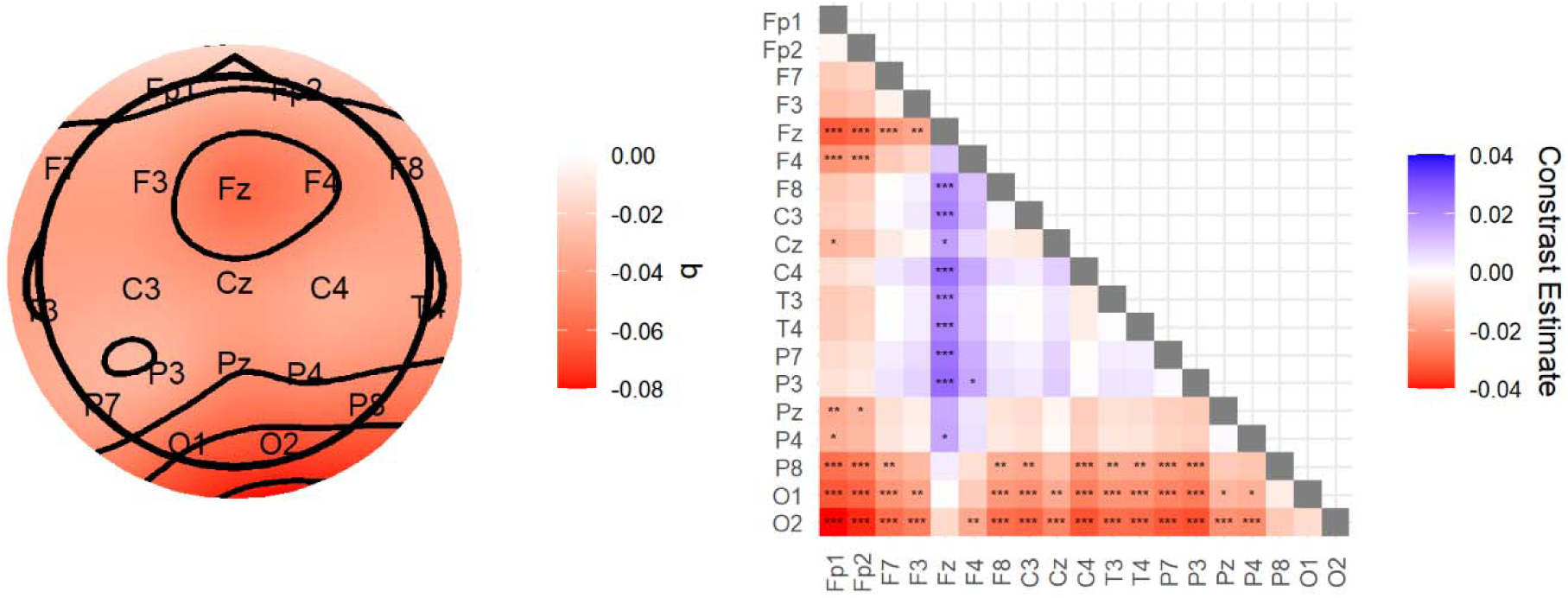
Two-way interaction between cortisol concentration and electrodes in predicting the exponent of the fitted 1/*f* slope. The left panel illustrates the scalp topographies of the estimated marginal slopes between cortisol concentration and exponent (*b*). The right panel illustrates the pairwise contrasts in *b* across electrodes. *** *p* < .001, ** *p* <. 01, * *p* < .05 (Holm-corrected).

Regarding the second hypothesis, there was a significant cortisol x resilience x electrode interaction effect, χ*^2^*(18, 54882.15) = 723.12, *p* < .001. As illustrated in Figure 2, simple slope analyses at ± 1 *SD* of trait resilience revealed a significant moderating effect, whereby the associations between cortisol concentration and 1/*f* exponent between low and high trait resilience differed at electrodes Fp1, Fp2, F4, F7, O1, and O2. Specifically, an increase in cortisol concentration was associated with a flatter 1/*f* exponent in individuals with low trait resilience, whereas an increase in cortisol concentration was associated with a steeper 1/*f* exponent in those with high trait resilience. By contrast, age (β = −0.006, *SE* = 0.005, *p* = .239), sex (β = −0.020, *SE* = 0.044, *p* = .658), and cortisol sampling time (β = −0.000, *SE* = 0.000, *p* = .332), were not significant predictors. However, epoch (β = 0.001, *SE* = 0.000, *p* < .001) was a significant predictor, indicating that the 1/*f* exponent increased in steepness over the baseline period.

**Figure 2.**
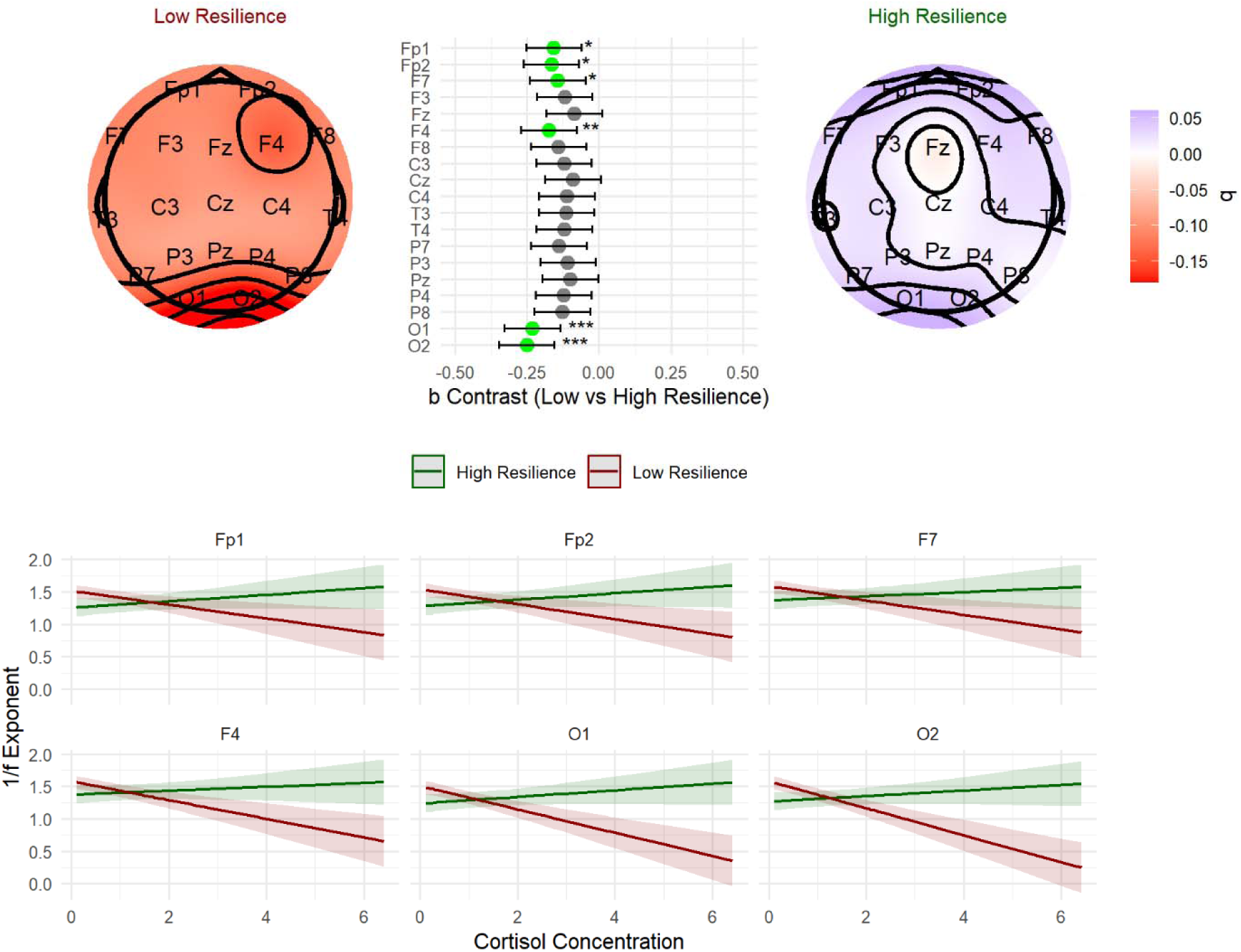
Three-way interaction between cortisol concentration, trait resilience, and electrodes in predicting the exponent of the fitted 1/*f* slope. The top left and right panels illustrate the scalp topographies of the simple slopes between cortisol and exponent (*b*) at low (−1 *SD*) and high (+1 *SD*) levels of trait resilience. The top centre panel illustrates the pairwise contrast in *b* between low and high levels of trait resilience at each electrode. Bottom panels depict simple slopes of cortisol predicting 1/*f* exponent separately at high and low trait resilience levels for electrodes showing significant contrasts. Error bars represent 95% confidence intervals. *** *p* < .001, ** *p* <. 01, * *p* < .05 (Holm-corrected).

**Figure 3.**
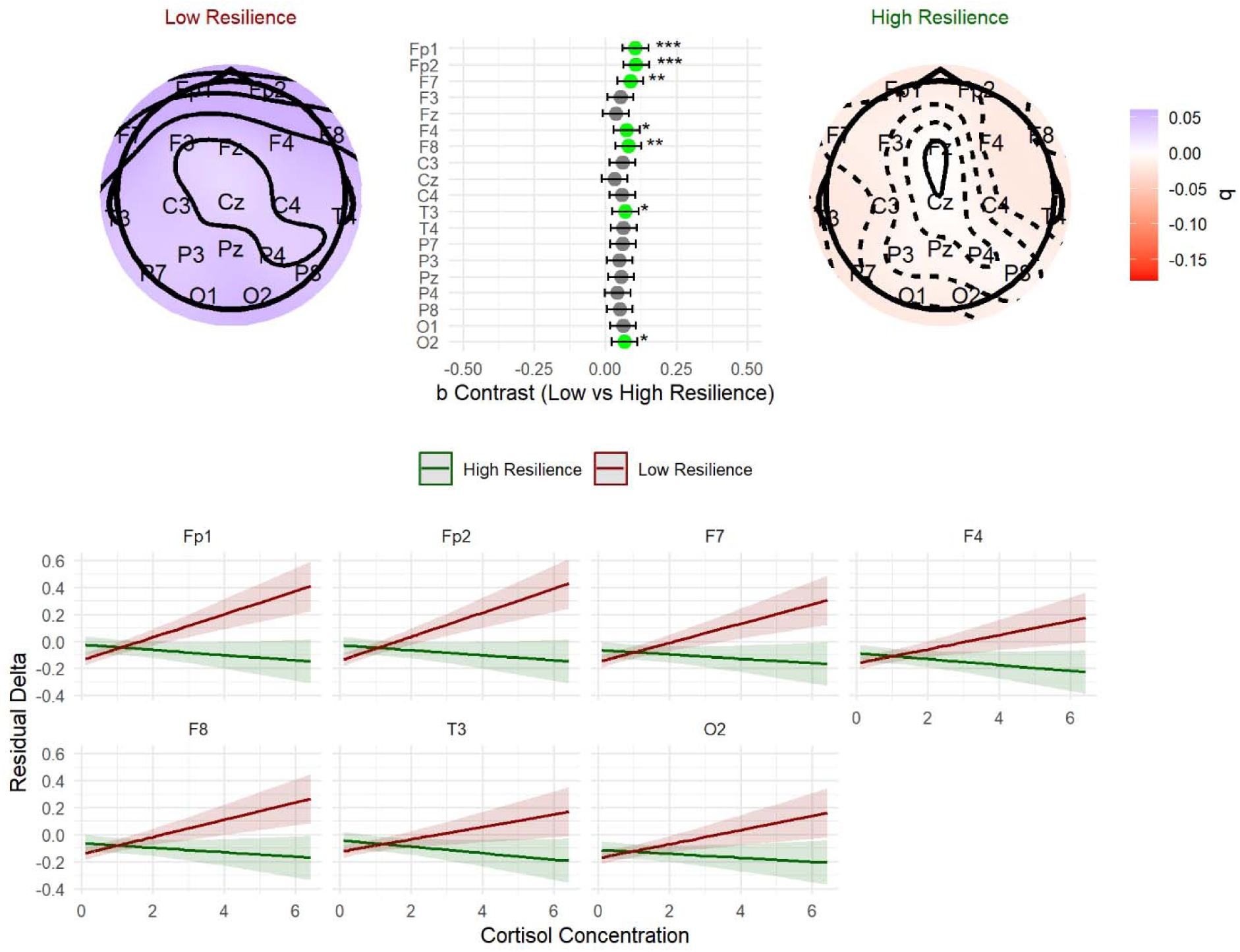
Three-way interaction between cortisol concentration, trait resilience, and electrodes in predicting residual delta activity. The top left and right panels illustrate the scalp topographies of the simple slopes between cortisol and exponent (*b*) at low (−1 *SD*) and high (+1 *SD*) levels of trait resilience. The top centre panel illustrates the pairwise contrast in *b* between low and high levels of trait resilience at each electrode. Bottom panels depict simple slopes of cortisol predicting residual delta activity separately at high and low trait resilience levels for electrodes showing significant contrasts. Error bars represent 95% confidence intervals. *** *p* < .001, ** *p* <. 01, * *p* < .05 (Holm-corrected).

### Specificity and Robustness Analyses of EEG Spectral Components

To establish the spectral specificity of our 1/*f* exponent findings, additional GAMs were conducted by replacing the outcome variable with alternative spectral components. The 1/*f* offset model, χ*^2^*(18, 54882.14) = 212.07, *p* < .001, delta model, χ*^2^*(18, 54882.15) = 719.99, *p* < .001, theta model, χ*^2^*(18, 54882.28) = 223.95, *p* < .001, alpha model, χ*^2^*(18, 54882.12) = 117.79, *p* < .001, and beta model, χ*^2^*(18, 54882.12) = 146.27, *p* < .001, revealed significant cortisol x resilience x electrode interaction effects. However, follow-up simple slope analyses did not identify electrode-specific moderation by trait resilience for the 1/*f* offset, theta, alpha, and beta models. By contrast, there was a significant moderating effect of trait resilience on the association between cortisol concentration and residual delta activity at electrodes Fp1, Fp2, F4, F7, F8, T3, and O2 (see Figure 3). Specifically, an increase in cortisol concentration was associated with enhanced delta activity in individuals with low trait resilience, whereas an increase in cortisol concentration was associated with a decrease in delta activity in those with high trait resilience.

**Figure 3.**
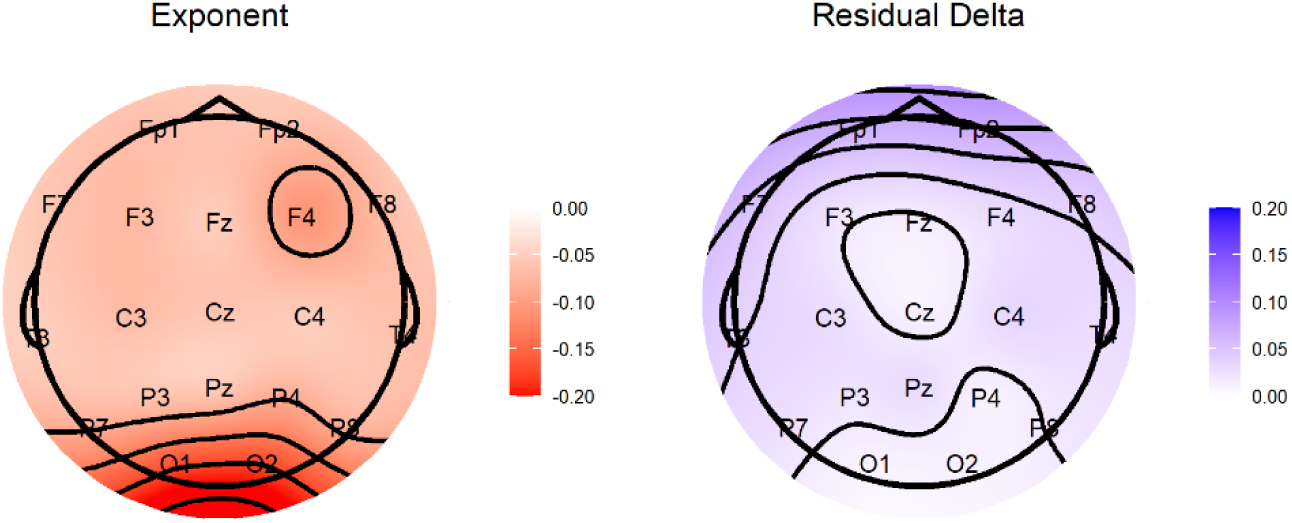
The topographies of the moderating effects of trait resilience on cortisol predicting 1/*f* exponent and residual delta activity. Note that the 1/*f* exponent was included as a covariate in the residual delta model, and residual delta activity was included as a covariate in the 1/*f* exponent model. The left and right panels depict scalp distributions of differences in cortisol-related simple slopes between low (−1 *SD*) and high (+1 *SD*) levels of trait resilience in predicting the two spectral components.

Given the spatial overlap between the moderating effects of trait resilience on the 1/*f* exponent and delta activity across prefrontal and occipital regions, we conducted additional spectral specificity analyses by alternatively including residual delta activity and 1/*f* exponent as covariates and outcome variables, respectively. The 1/*f* exponent model revealed a significant cortisol x resilience x electrode interaction effect, χ*^2^*(18, 54881.14) = 594.60, *p* < .001, even after controlling for the effects of delta activity. Post hoc simple slope analyses revealed that only the effects at electrodes O1 (*p* = .004) and O2 (*p* = .002) remained. The delta model also revealed a significant cortisol x resilience x electrode interaction effect, χ*^2^*(18, 54881.13) = 804.95, *p* < .001, even after controlling for the effects of 1/*f* exponent. However, only the effects at electrodes Fp1 (*p* = .023) and Fp2 (*p* = .010) remained. Overall, the additional spatial specificity analyses revealed a clear regional dissociation (see Figure 3). Specifically, the moderating effect of trait resilience on delta activity remained pronounced only in prefrontal scalp regions, whereas the effects on the 1/*f* exponent were largely localised to occipital scalp regions.

### Specificity and Robustness Analyses of Psychosocial Moderating Effects

To establish the psychosocial specificity of the moderating effects of trait resilience on 1/*f* exponent and delta activity, additional GAMs were conducted by replacing trait resilience with psychological well-being and sleep quality as alternative moderators. The psychological well-being models revealed significant three-way interaction on the 1/*f* exponent, χ*^2^*(18, 54881.14) = 181.79, *p* < .001, and delta activity, χ*^2^*(18, 54881.12) = 39.31, *p* = .003. In addition, the sleep quality models also revealed significant three-way interaction on the 1/*f* exponent, χ*^2^*(18, 54502.14) = 104.57, *p* < .001, and delta activity, χ*^2^*(18, 54502.12) = 243.97, *p* < .001. However, subsequent simple slope analyses did not identify significant electrode-specific moderation by psychological well-being or sleep quality for either spectral component. To further investigate the robustness of the observed moderating effect of trait resilience, both psychological well-being and sleep quality were included as covariates. The 1/*f* exponent model revealed a significant interaction effect, χ*^2^*(18, 54502.14) = 600.24, *p* < .001. Post hoc simple slope analyses showed a consistency with prior patterns, such that the moderating effect were localised at electrodes O1 (*p* = .003) and O2 (*p* = .002). The delta model also revealed a significant interaction effect, χ*^2^*(18, 54502.12) = 826.99, *p* < .001, with post hoc simple slope analyses showing the moderating effects localised at electrodes Fp1 (*p* = .021) and Fp2 (*p* = .010), which was consistent with prior patterns.

## Discussion

In the present study, we examined individual differences in the association between cortisol and aperiodic neural activity with trait resilience as a potential moderator. Our results revealed that resting cortisol concentration was associated with 1/*f* exponent, particularly over the occipital, parietal, and frontocentral scalp regions. However, this relationship was qualified by varying levels of trait resilience. Specifically, an increase in cortisol concentration was associated with a flatter 1/*f* slope in individuals with low trait resilience. Conversely, the association was reversed, such that increased cortisol concentration was associated with a steeper 1/*f* slope in those with high trait resilience. Topographically, this moderating effect of trait resilience was most pronounced across prefrontal and occipital scalp regions. However, trait resilience also showed similar spatial patterns of moderating effect on delta activity. Subsequent spectral specificity analyses revealed that the moderating effect on aperiodic activity remained robust only in occipital regions after controlling for delta activity. Conversely, regarding delta activity, the effects remained only in the prefrontal regions after controlling for aperiodic activity.

Together, our findings indicate that trait resilience moderates how individual differences in cortisol levels is associated with both aperiodic and delta activity, with these effects showing spatially dissociable patterns localised to occipital and prefrontal regions, respectively.

Drawing on the neurophysiological functional interpretations of the 1/*f* slope (R. Gao et al., 2017; Voytek et al., 2015), our findings pertaining to aperiodic activity suggest that increased cortisol in individuals with lower trait resilience shifts the excitatory-inhibitory balance toward greater excitation or increases neural noise in occipital regions. In contrast, increased cortisol in those with higher trait resilience appears to shift this balance toward greater inhibition or decrease neural noise. One plausible explanation is that highly resilient individuals better maintain neurophysiological homeostasis despite increased neuroendocrine activity. Given that a flatter 1/*f* slope is associated with increased physiological arousal (Colombo et al., 2019; Lendner et al., 2020; Pfeffer et al., 2022), it appears that individuals with lower trait resilience remain in a state of heightened cortical excitation when cortisol is elevated, reflecting increased physiological arousal even during rest. In contrast, individuals with higher trait resilience may be better able to attenuate the influence of elevated cortisol during rest, thereby maintaining a lower level of arousal state characterised by reduced cortical excitation. Importantly, the occipital localisation of the observed moderating effect may point to a potential pathway involving reduced sensitivity of posterior sensory cortical systems to cortisol-related arousal, which future research should investigate further.

Regarding delta activity, the association between cortisol concentration and prefrontal delta activity also differed as a function of trait resilience. Specifically, amongst individuals with lower trait resilience, higher cortisol concentrations were associated with increased prefrontal delta activity, whereas the opposite pattern was observed amongst individuals with higher trait resilience. One possible interpretation of this differential association is that trait resilience may reflect neural compensatory processes that modulate the cortical effects of elevated cortisol. That is, individuals with higher trait resilience may be more effective at regulating cortisol-related influences on cortical dynamics, thereby maintaining more adaptive functioning under conditions of elevated physiological stress. This interpretation is broadly consistent with prior conceptualisations of resilience as an active neurobiological process involving dynamic behavioural, neural, molecular, and hormonal adaptations to stress (Russo et al., 2012).

Supporting this account, neuroimaging studies have shown that highly resilient individuals exhibit stronger top-down inhibitory control from prefrontal regions over the amygdala (Bolsinger et al., 2018), which is a key excitatory regulator of the HPA axis (Herman et al., 2005). In addition, prefrontal neural regions are also known to play a central role in the generation and regulation of delta oscillatory activity, with enhanced delta activity during wakefulness reflecting disrupted or suboptimal homeostatic functioning (Knyazev, 2007, 2012). Hence, the localisation of the delta activity findings may reflect the engagement of prefrontal regulatory systems that serve to buffer against the effects of elevated cortisol. Increased prefrontal delta activity in individuals with lower trait resilience may therefore reflect a dysregulated top-down inhibitory process, potentially indicating reduced capacity for adaptive homeostatic regulation.

Overall, the present findings indicate that HPA-axis functioning is differentially associated with both aperiodic and low-frequency neural dynamics as a function of trait resilience. Rather than reflecting a uniform cortisol-cortical relationship, cortisol-related neural effects appear to be systematically shaped by individual differences in the capacity to cope with stress or adversity, with distinct spatial and spectral signatures across the scalp. This pattern of findings was not better explained by orthogonal psychosocial constructs, such as the previous night’s sleep quality and psychological well-being. Extending recent work demonstrating that individuals with a flatter 1/*f* slope were at higher risk of depression recurrence when under high stress levels (Schantell et al., 2026), our neuroendocrinological findings further suggest this psychopathological vulnerability may be conditionally depend on trait resilience via its modulating effect on cortisol-cortical dynamics. Hence, our study highlights the importance of considering individual differences in trait resilience when examining spontaneous cortisol-cortical relationship. In addition, our study also raises the possibility that the combination of aperiodic and low-frequency oscillatory activity may serve as a candidate neurophysiological signature of trait resilience.

There are several limitations pertaining to our study that should be noted. First, our analyses relied on a single salivary cortisol measure, limiting the ability to accurately characterise diurnal patterns in cortisol secretion. Although cortisol sampling time was included as a covariate to partially account for the time-of-day effect, this approach cannot fully substitute for repeated measurements taken multiple times within and across days. Future research would therefore benefit from repeated sampling across multiple time points, including awakening and diurnal decline phases, to more accurately capture individual differences in HPA axis functioning.

Second, while the spatial distribution of the effects across the scalp suggests potential functional differentiation across cortical regions, the cross-sectional design precludes any inference regarding directionality cortisol-cortical dynamic. Future research should employ orthogonal manipulations combining pharmacological approaches (e.g., hydrocortisone administration) with cognitive control tasks to disentangle whether the observed prefrontal and occipital effects genuinely reflect prefrontal regulatory processes and posterior cortical sensitivity, respectively.

## Contributions

K.F.A.L. & T.M.C.L., conceptualisation, supervision, and project management, K.F.A.L., L.L., & T.M.C.L., methodology and resources, K.F.A.L., L.L., & T.A.S., investigation and data curation, K.F.A.L., formal analysis, software, validation, and visualisation, K.F.A.L., & T.A.S., writing-original draft, K.F.A.L., L.L., T.A.S., & T.M.C.L., writing-review & editing, T.M.C.L., funding acquisition. All authors contributed to the critical revision of the manuscript and approved the final version for submission. ^#^ These authors contributed equally.

## Data and Code Availability

The data, while not publicly available due to participant privacy, are available from the corresponding author upon reasonable request. All code has been deposited at (https://github.com/alvinleekarfye/alvin-aperiodic-cortisol-restinganalysis-2026) and is publicly available at (https://doi.org/10.5281/zenodo.21274239).

